# Life-history evolution under artificial selection in a clonal plant

**DOI:** 10.1101/2025.10.01.679824

**Authors:** Christina Steinecke, Isabeau Lewis, Jeremiah Lee, Jannice Friedman

## Abstract

The response of natural populations to selection and the role of genetic correlations in constraining or facilitating evolutionary change is fundamental to adaptation. We use artificial selection to investigate the evolutionary response of clonal reproduction in the common monkeyflower (*Mimulus guttatus*), a species with extensive life history variation. We first characterize the standing genetic variation in a single perennial population, and then conduct four generations of divergent artificial selection on stolon number—the mechanism of clonal reproduction in this species. To start, stolon number had moderate heritability (*H*²=0.25) and was negatively genetically correlated with reproductive traits. Artificial selection produced a clear but asymmetrical response. High selection lines made significantly more stolons, while low lines diverged less from controls. Analyses of ***G*** matrices revealed that selection not only changed trait means but also genetic correlations, with high lines diverging more in multivariate genetic architecture. Our results demonstrate that single populations harbor sufficient genetic variation to respond rapidly to selection on clonality, and the response is shaped by existing patterns of genetic covariation. The capacity for rapid evolution of clonal traits is particularly relevant as climate change alters selection and shifts the relative advantages of sexual versus clonal life-history strategies.

## INTRODUCTION

The capacity of natural populations to respond to changing environments depends on the standing genetic variation of traits under selection and the strength and direction of selection (Lande & Arnold, 1983). Furthermore, the rate and direction of trait evolution may be either promoted or constrained by selection acting on genetically correlated traits (Antonovics, 1976; Blows and Hoffmann, 2005), which show a correlated response in proportion to the strength of selection and the magnitude of the genetic correlation (Lande, 1979; Arnold, 1992; Schluter, 1996). Demonstrating natural selection can be challenging, and artificial selection can represent a tool for testing the evolutionary response to selection and for exploring potential constraints on adaptive evolution in response to altered selection (Schluter, 1988; Conner, 2003; Brakefield, 2003), at least in terms of short-term evolutionary responses of populations.

Evolutionary outcomes depend not only on the direction and strength of selection but also on the genetic (co)variances among traits (Lande, 1979; Lande and Arnold, 1983). Indeed, evolutionary paths may follow genetic lines of least resistance (Schluter, 1996) that reflect the multivariate trait combination with the greatest amount of genetic variation. Additionally, environmental variation that favours particular combinations of traits can lead to the evolution of genetically correlated trait combinations (e.g., Rodd and Reznick, 1997; Walter et al., 2024). Thus, it is not always clear what traits are the target of selection, as combinations of traits often evolve in a coordinated way. Understanding the relationship between phenotypic divergence among populations and genetic variation and local selection within populations remains a major goal of evolutionary biology.

Adaptive divergence in life-history traits represent classic examples of multivariate phenotypes that result from the joint action of genetic correlations and selective conditions (Wadgymar et al., 2022). Life-history strategies generally encompass traits that determine key life cycle parameters like timing and investment in reproduction, growth, and survival. Life-history theory predicts the existence of trade- offs between different fitness components, that could arise due to genetic correlations (e.g., QTL affecting growth and flowering in *Arabidopsis thaliana*; Mitchell-Olds, 1996) or through allocation trade-offs due to finite resources (e.g., negative phenotypic correlations between early fecundity and adult lifespan in *Drosophila melanogaster*; Flatt, 2005; Flatt and Schmidt, 2009). The concept of trade- offs and differential allocation of limiting resources to reproduction versus growth and maintenance for survival is central to life-history theory.

Flowering plants display enormous variation in life-history strategies, including the ability to reproduce both sexually and asexually. Approximately 80% of plant species can reproduce clonally via vegetative propagation (Fryxell, 1957; Klimeš et al., 1997). Clonal reproduction can be beneficial in environments unfavourable for sexual reproduction (e.g., when mates are limiting), and can facilitate the guaranteed and quick production of identical genotypes and the transmission of genes to the next generation (Roughgarden, 1991). Clonality also increases opportunities to forage for resources and spreads the risk of death among ramets (Vallejo-Marín et al., 2010; Wildová et al., 2007). Clonal structures can influence how a plant grows in both time and space, affecting mating outcomes (Vallejo-Marín et al., 2010) and shape patterns of genetic and phenotypic variation (Harper, 1977; Leme da Cunha et al., 2022). Selection may act on clonal traits if they increase survivorship and reproductive value of clonal offspring (Fisher, 1930; Solbrig and Simpson, 1977). Sexual reproduction, in contrast, generates genetic diversity that can allow for adaptation to changing environments as well as reduce the accumulation of deleterious mutations (Fisher, 1930; Otto and Lenormand, 2002; Agrawal, 2006).

Although possessing both clonal and sexual reproductive modes has clear advantages for perennial species (Silander, 1985), these two activities can also interfere with one another, resulting in various forms of antagonism including potential trade-offs between clonal and sexual reproduction.

The common monkeyflower (*Mimulus guttatus* DC; syn. *Erythranthe guttata* DC GL Nesom; Phrymaceae) is a useful species to examine the effects of selection on clonal and sexual reproduction. The species is a predominantly outcrossing herbaceous angiosperm that displays extensive variation in sexual and clonal reproduction (Hall and Willis, 2006; van Kleunen, 2007; Friedman et al., 2015). It reproduces sexually with bee-pollinated, hermaphroditic flowers and clonally with stolons, which are horizontal vegetative branches that grow from axillary meristems and root at each node (Vickery, 1978; van Kleunen, 2007; Coughlan et al., 2021). Throughout much of its range in western North America, derived annual populations occur in dry, transient habitats and ancestral perennial ecotypes occur in areas with persistent soil moisture (Hitchcock and Cronquist, 1973; Vickery, 1978). Stolons are key to the vegetative (perennial) strategy in *M. guttatus,* as the following season’s rosettes form along their nodes (Friedman et al., 2015), and populations vary in their stolon traits (Nesom, 2012; Friedman, 2015; Coughlan et al., 2021). Annuals do not produce stolons in nature (Vickery, 1978). Additionally, plants that flower earlier tend to make fewer stolons and vice versa; a pattern that appears to be due to pleiotropic QTL in between-population crosses (Hall et al., 2006; Friedman et al., 2015), and within- population genetic correlations show the same association (Rubin et al., 2018). Compared to flowering traits, research on stolons is limited and it is unclear how much variation exists within populations and how selection on clonal reproduction might affect phenotypic and genetic variation.

To evaluate the capacity for populations to respond to selection on clonality, and the influence on correlated multivariate traits, we applied divergent artificial selection on stolon production. We began by choosing a perennial population that was roughly intermediate in its stolon traits in relation to other perennial *Mimulus guttatus* (Friedman et al., 2015). We used a set of open-pollinated seed families and grew plants in a common greenhouse environment to determine the standing genetic variation and heritability of a range of phenotypic traits. In this founding population, we also tested for genetic correlations between traits to establish the multivariate phenotype that may respond to selection on stolon number. We then applied four generations of divergent artificial selection on stolon number by randomly mating only the individuals that produced the greatest 50% (2 replicate groups), the lowest 50% (two replicate groups), or randomly mated (control). In the final generation, in addition to the selection and control lines, we grew individuals from the original parental population to evaluate the evolutionary response to selection, as well as to detect any unintended consequences such as changes in trait means or variance unrelated to direct selection. We expected selection lines to differ significantly from the control and ancestral populations in stolon number, and potentially in correlated traits if selection also acts on trait covariations. We further tested whether the multivariate phenotype and genetic variance-covariance (***G***) matrix changed through time, and whether divergent selection on stolon number resulted in contrasting suites of traits. This allows us to better understand how selection on one trait can shape the evolution of correlated traits and the overall genetic architecture within a population. Ultimately, we are interested in examining the ability of perennial populations to respond to selection on clonality, but also to understand more generally how selection on a key life-history trait may affect the ability for populations to adapt to different or changing environments.

## METHODS

### Establishment of the Base Generation

We used open-pollinated seed families collected in 2013 from a single perennial population (GWD), located near Sequoia National Forest, California (N35°57’56", W118°28’42"). We germinated up to 50 seeds from 35 seed families on Sunshine® Mix #1 (Sun Gro Horticulture; www.sungro.com) potting soil in December of 2020. Seeds were planted in 6.35-cm pots and stratified at 4°C in the dark for three days. We placed the pots into a greenhouse at the Queen’s University Phytotron with conditions set at 15.5-h photoperiod, 21/18°C day/night. Germination in this species is very high (>95%) and highly synchronous (within 1-3 days). Once seedlings had open cotyledons, we haphazardly transplanted 24 seedlings from each family (except for one family with 14 seedlings due to low germination; N = 836) into individual 6.35-cm pots. We placed plants into a random block design and rotated them weekly to limit effects from position in the greenhouse. All plants were bottom-watered for one hour daily and fertilized with 240ppm of 10N-10P-10K weekly.

Five weeks after germination we counted the number of stolons and measured the length of the longest stolon on each individual. Nine weeks after germination, we again counted the number of stolons and measured the length of the longest stolon, and measured stem thickness between first and second nodes, the number of flowers, number of nodes, and number of branches on each individual.

Additionally, we measured the number of days from germination to first flowering, as well as the length and width of the first flower. Although plants are perennial, their growth slows, their leaves senesce, and they stop producing new flowers after about 9 weeks in the greenhouse (in nature, their above-ground material senesces at the end of summer).

Within each of the 35 families, siblings were split equally into two groups (N = 12 of 24 sibs per group) to act either as maternal or paternal parents. We used a factorial crossing design to create 395 unique pairs with the condition that individuals were paired across families to maximize different between-family crosses, and ensured all families were equally represented as maternal and paternal parents. We performed unidirectional hand pollinations to create outbred full-sibling seed – we tapped pollen from anthers of the assigned paternal parent onto a glass microscope slide and used a toothpick to place the pollen on the stigmatic lobe of the assigned maternal parent and marked the flower with a small piece of tape. We collected ripe seed pods in coin envelopes and stored them at room temperature until seeds were sown for the next generation.

### Artificial Selection Regime

We began the first generation of selection with seed from 274 of the 395 unique family pairs created from the base generation. To maximize and evenly represent genetic variation, we used approximately eight crosses from each of the original 36 maternal families. We germinated five seeds per full-sib family in April of 2021 into a single 6.35-cm pot and followed the same stratifying, growing, watering, and fertilizing conditions as outlined above. We transplanted each of the five seedlings into individual 6.35-cm pot (N = 1370). We placed one of each sibling into each of five groups: two replicate high (**H1** and **H2**), two replicate low (**L1** and **L2**), and one control (**C**), so that there were N= 274 plants per treatment (Figure S1). We distributed plants into a randomized block design and rotated them weekly.

Artificial selection was imposed at five weeks after germination, following the measurement of stolon number and length. We ranked individuals by their stolon number and retained individuals from the top (**H**) or bottom (**L**) 50% of the distribution within each group to use as parents for the next generation (Figure S1). When multiple individuals tied in the ranking, we ranked them by stolon length and retained the individuals with the longest (**H**) or shortest (**L**) stolons, respectively. In the first generation only, plants that did not flower or whose flowering did not overlap with other plants within their selection line, were replaced with an unselected sibling (N = ∼15). Within each selection group we mated the selected individuals at random, and within the control group we randomly crossed all individuals, with the proviso that we did not mate full siblings. We collected ripe seed pods into coin envelopes and stored them at room temperature before using them for the next generation.

To maintain a constant population size, for the next three generations of selection we began with four full siblings from each family within each selection/control line (Figure S1). Seeds for each generation were sown in July of 2021 (generation 2), October of 2021 (generation 3), and March of 2022 (generation 4). Growth conditions, measurements of traits, and the selection regime were identical to the first generation of selection outlined above, with one exception. Between the first and second generation of selection, the halogen lighting in the greenhouse was upgraded to full spectrum LED lighting due to scheduled facility upgrades. Because selection generations were grown sequentially, any environmental variance between generations had the potential to obscure the response to selection by increasing the environmental variance component of heritability (i.e., parents and offspring do not experience identical environments). To account for this, in the final generation, we grew two full siblings from families created in the base generation (i.e., full siblings of first-generation individuals and ancestors of final-generation individuals, hereby referred to as the Ancestral group). This also allowed us to test for effects of genetic drift and/or unanticipated selection by comparing the ancestral group to the control group.

## Statistical Analyses

### Standing Genetic and Phenotypic Variation

We performed all analyses in R version 4.2.0 (R Core Team, 2022). We began by estimating the distributions, heritabilities, and variance components of the 11 traits in the base generation using linear mixed model analyses fitted with the lmer function from the ‘lme4’ package (version 1.1-30; Bates et al., 2015). Each model included family as a random effect and block as a fixed effect. Variance components for the random effects were estimated using restricted maximum likelihood (REML). To assess the significance of the family random effect, we conducted likelihood ratio tests comparing model fit with and without the random effect using the ‘lmerTest’ package (Kuznetsova et al., 2017). We extracted Best Linear Unbiased Predictions (BLUPs) for each family, variance components, and calculated population- and family-level means and standard deviations for each trait. To assess genetic relationships among traits, we performed pairwise regressions using family-level BLUPs, with a select subset of focal traits representing key aspects of clonal and sexual reproduction: stolon number, flowering time, flower number, and corolla width (flower size). These traits were chosen based on their ecological relevance to vegetative propagation and reproductive allocation in *M. guttatus* (Hall and Willis, 2006; Lowry and Willis, 2010; Friedman et al., 2015).

Using the variance components in the above models, we estimated genetic variance (***V_G_***) of all traits as the inverse of the expected relatedness of sibling offspring (Lynch and Walsh, 1998). Because the base generation used open-pollinated seed, we assumed that offspring within each family were half-siblings with different pollen donors, and estimated ***V_G_*** as four times the family variance component.

Total phenotypic variance (***V_P_***) was calculated as the sum of the residual variance (***V_E_***) plus the genetic variance (***V_G_***) as estimated above. Finally, we calculated broad-sense heritability (***H^2^***) as the ratio of the estimated genetic variance to the total phenotypic variance (Lynch and Walsh, 1998).

We used several complementary approaches to investigate multivariate phenotypic variation. First, we performed a Principal Component Analysis (PCA) on the 11 phenotypic traits to identify the primary axes of variation and the loadings of each trait. We summarized the strength and direction of pairwise trait associations using a Pearson phenotypic correlation matrix. Finally, we estimated the genetic variance-covariance (***G***) matrix, using Bayesian generalized linear mixed-effect models with the MCMCglmm function from the ‘MCMCglmm’ package (Hadfield, 2010). To enable comparative analyses with the selection lines (see below), we assigned each plant in the base generation a unique paternal parent identifier, since pollen donor identity was unknown in this generation. This results in a possible under-estimate of the G-matrix values if some individuals actually share pollen donors. Our priors assumed that maternal- and paternal-family covariance matrices were equal to 0.25 × the phenotypic covariance matrix (***P***), with nu = *n* + 1, with *n* as the number of traits (McGoey & Stinchcombe, 2021). We ran full models with both maternal family and paternal family as random effects and block as a fixed effect for 500,000 iterations, with a burn-in of 5,000 and a thinning interval of 500, resulting in 1,000 iterations sampled from the posterior distribution.

Genetic correlations (***r_g_***) between traits were calculated by standardizing the off-diagonal covariance components of the ***G*** matrix. Specifically, each covariance was divided by the square root of the product of the corresponding trait variances, allowing interpretation of the strength and direction of genetic associations between traits on a common scale.

### Univariate Response to Selection

To assess the direction and magnitude of change in stolon number across generations and selection lines, we used linear mixed models with a Poisson error distribution. Stolon number was the dependent variable, with generation × selection line as a fixed effect interaction, block (nested within generation) as a fixed effect, and family as a random effect to account for shared genetic background. We then compared pairwise estimates of marginal means (EMMs) for stolon number among all selection and control lines within each generation using the ‘emmeans’ package (Lenth, 2022). In total, we performed 10 pairwise contrasts per trait per generation, with error rates adjusted using Tukey’s method for multiple comparisons.

We calculated the selection differential (***S***) within each generation and selection line as the difference in the mean number of stolons in the population (X^-^) and the mean number of stolons of the selected parents (***X_P_***); the selection response (***R***) as the difference in the mean number of stolons in the population and the offspring of the selected parents (***X’***); and estimated narrow-sense heritability (***h*^2^**) from the breeder’s equation (i.e., ***h*^2^ = *R/S***; Falconer and Mackay, 1996). Standard errors for each estimate of ***h*^2^** were calculated following methods outlined in Burgess et al. (2007) and Roff (1997) to account for variance that may be due to genetic drift.

Because high and low lines were grown simultaneously, within-generation phenotypic differences between them are due to genetic change in both directions (Conner et al., 2011). However, between-generation differences in growth conditions could inflate the environmental variance. We therefore used linear mixed models to compare stolon number at week five (when selection was imposed) of the entire first generation, the control group in the final generation, and the ancestral population, with block as a fixed effect nested within generation and family as a random effect.

Significant differences between groups could indicate that the response to selection was impacted by between-generation differences in growth conditions and/or the occurrence of unintended selection.

### Multivariate Response to Selection

Using the same methods outlined above, we compared pairwise EMMs for the same subset of traits as the base generation (flowering time, flower number, and corolla width), performed a PCA, and estimated the ***G*** matrix for each selection line after selection, as well as for the ancestral population.

Before conducting ***G*** matrix comparative analyses, we first assessed whether the observed structures in the ***G*** matrices reflected genuine genetic variance-covariances, or whether they were an artifact of sampling error or environmental noise. We followed previously established methods (Walter et al., 2018; McGoey and Stinchcombe, 2021; Henry and Stinchcombe, 2023), and randomly assigned families (without replacement) within each group (G0, the ancestral population, and within each selection line after selection), and re-estimated their ***G*** matrices. We iterated this randomization and ***G*** matrix estimation 1000 times per group, creating seven independent null distributions of ***G*** matrices representing the expected genetic (co)variance structure under random sampling alone. For each group, we calculated the Frobenius distance between the observed ***G*** matrix and every matrix in its corresponding null distribution to quantify multivariate divergence. We visualized these null distributions alongside the observed value using histograms, with 95% quantile intervals indicated to assess significance (Figure S2; Table S1). The observed ***G*** matrices for all groups differed significantly from their respective null distributions, confirming that the observed genetic structures were not explained by sampling alone (Figure S2; Table S1) and justifying further analyses.

To assess whether ***G*** matrices changed over time and due to selection, we compared the ancestral population to the control and selection lines using a number of techniques described in Henry and Stinchcombe (2023), including ***G*** matrix size comparisons, eigenvalue comparisons, and Krzanowski’s common subspace analysis. We began by bootstrapping ***G*** matrix distributions for each group by randomly subsampling 100 individuals with replacement and constructing a ***G*** matrix as described above, repeating this process 1000 per times per group (n = 6 bootstrap distributions). These distributions accounted for sampling error and provided estimates of variability in key summary statistics. We quantified ***G*** matrix size, representing the total amount of quantitative genetic variation, by calculating the trace (sum of the eigenvalues) for each ***G*** matrix and compared the resulting distributions of ***G*** matrix sizes among groups using an Analysis of Variance.

We used Krzanowski’s common subspace analysis to determine whether the bootstrapped distributions of the ***G*** matrices exhibited similar patterns of genetic variation. This method identifies subspaces of dimensional space that represent the most significant directions of variation (Krzanowski, 1979). If the subspaces with the most genetic variation are similar across groups, it suggests that genetic structures (e.g., covariances due to linkage disequilibrium or pleiotropy) are shared (McGooey and Stinchcombe, 2021; Henry and Stinchcombe, 2023). For each ***G*** matrix, we calculated its eigenstructures, identified the principal components and their corresponding eigenvectors that explained maximum genetic variance, and extracted the subspaces associated with the leading eigenvalues. These subspaces were then flattened into data frames consisting of their corresponding eigenvectors and compared across groups using an Analysis of Variance.

## RESULTS

### Standing Genetic and Phenotypic Variation

All traits measured in the base generation showed significant genetic variation, assessed by comparing mixed models with and without a random effect of family (Table 1). Estimates of broad-sense heritability ranged from 0.17 to 0.48, with stolon traits approximately intermediate relative to other traits (week 5: ***H²*** = 0.25; week 9: ***H²*** = 0.22; Table 1). The number of stolons ranged from 0 to 8 stolons (mean = 3.08) at week 5 and from 2 to 15 stolons (mean = 7.6) at week 9 (Table 1; Figure S3). Patterns of multivariate phenotypic variation in the base generation were evident from the phenotypic PCA, pairwise Pearson correlation matrix, and genetic variance-covariance (***G)*** matrix (Figure 1A-D; Table 2; Table S2; Table S3; Table S4). The first principal component (PC1) accounted for 28.7% of the variation, with all traits having positive loadings (Table 2; except flowering time which is traditionally coded orthogonally to other traits). This indicates that PC1 reflects overall plant size. On PC2 (explaining 15.8% of variation), there was a clear divergence between clonal traits and flowering traits (Figure 1A). Stolon traits such as number and length had negative loadings, while floral traits such as flower number, branch number, and nodes, had strong positive loadings on this axis (Table 2). Both phenotypic (ρ) and genetic correlations (***r_g_***; calculated by standardizing the off-diagonal covariance components of the ***G*** matrix) revealed similar trait associations, indicating a genetic trade-off between clonal and reproductive investment (Table S2; Table S4). Regressions on family-level BLUPs indicated that at the initiation of flowering (i.e., measured at week 5), individuals producing more stolons also grew longer stolons (ρ = 0.63, *P* < 0.001; ***r_g_***= 0.22; Table 2; Table S2; Table S4). By the end of the experiment (i.e., measured at week 9) plants that produced more stolons tended to flower later (ρ = 0.38, *P* < 0.001; ***r_g_*** = 0.15; R^2^ = 0.24, *P* = 0.002; Figure 1B), produce fewer flowers (ρ = 0.41, *P* < 0.001; ***r_g_*** = 0.17; R^2^ = 0.18, *P* = 0.006; Figure 1C), and produced smaller flowers (ρ = -0.23, *P* < 0.001; ***r_g_*** = -0.24; R^2^ = 0.02, *P* = 0.19; Figure 1D) (Figure 1A; Table S2; Table S4), although for the latter the genetic correlation was not statistically significant.

**Figure 1.**
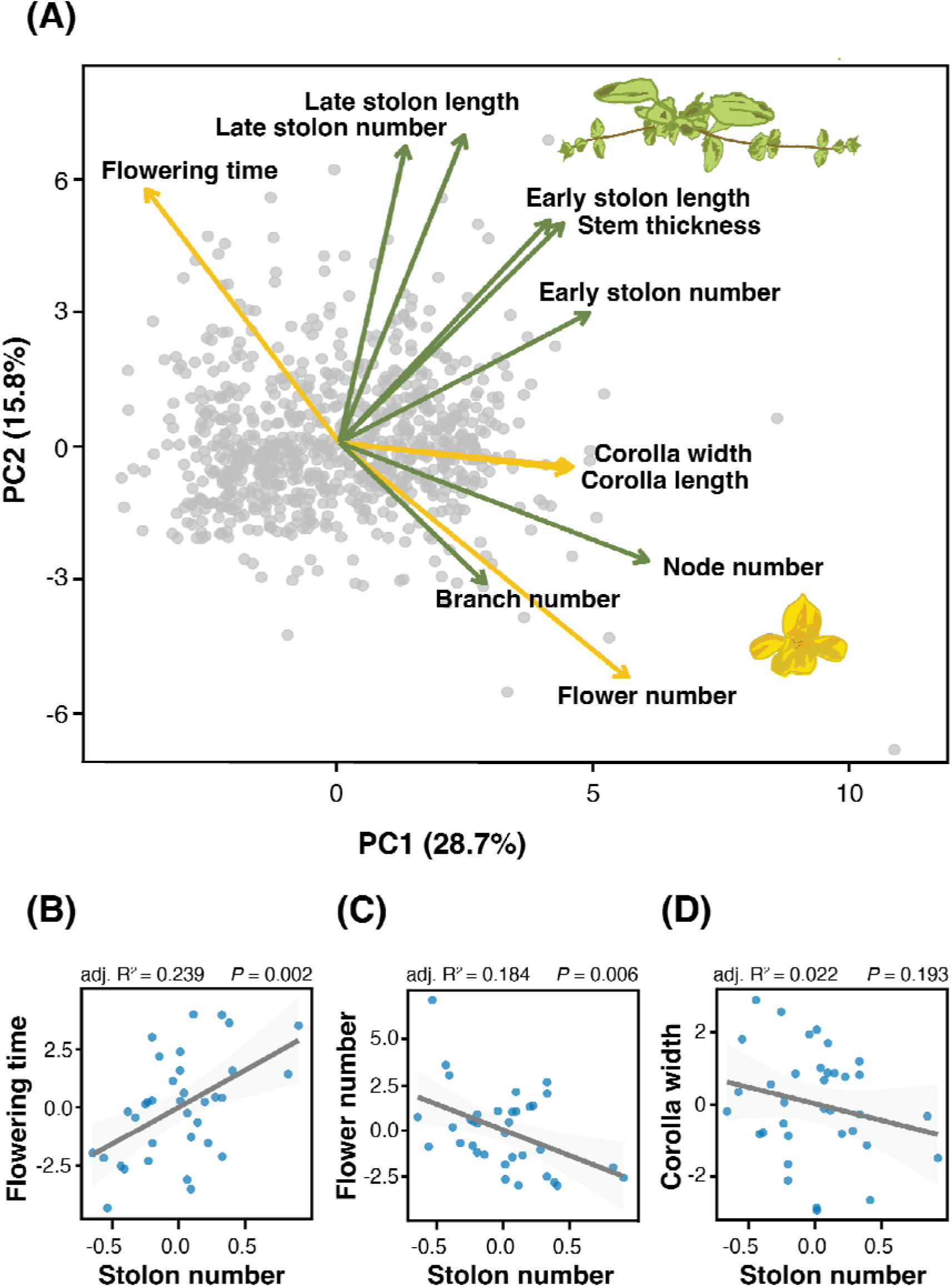
Standing phenotypic variation in traits measured in the base generation of *Mimulus guttatus*. (A) Principal Component Analysis (PCA) of phenotypic variation in the base generation. Points represent individual plants, and arrows indicate the loadings (direction and strength) of each phenotypic trait on the first two principal components. (B-D) Linear regressions between stolon number and floral traits (B: flowering time, C: flower number. D: corolla width) in the base generation, where each point represents an individual plant’s Best Linear Unbiased Predictor (BLUP), calculated from a mixed-effects model that included family as a random effect and experimental block as a fixed effect. Adjusted R^2^ and *P*-values are displayed in each panel to indicate the strength and significance of the pairwise trait associations in the base population.

**Table 1.** Means and standard deviations for phenotypic traits of *Mimulus guttatus* in the base generation (outbred field collected seed) grown in the greenhouse. Estimates of genetic variance (***V_G_***), phenotypic variance (***V_P_***), and broad-sense heritability (***H^2^***) are calculated from the variances associated with the random effect of family in linear mixed models. The significance of the random effect of family was assessed using likelihood ratio tests comparing models with and without it, and the Chi-squared statistic χ^2^, df=1) and corresponding *P*-values are reported.

| Trait | Mean + SD | $V_G$ | $V_P$ | $H^2$ | $\chi^2$ | $P$-value |
| --- | --- | --- | --- | --- | --- | --- |
| **Week 5** |  |  |  |  |  |  |
| Stolon number | 3.08 ± 1.16 | 0.45 | 1.84 | 0.25 | 26.13 | < **0.001** |
| Stolon length (mm) | 6.26 ± 3.02 | 39.13 | 85.53 | 0.46 | 85.40 | < **0.001** |
| Corolla length (mm) | 34.96 ± 2.12 | 4.81 | 28.32 | 0.17 | 11.51 | **0.0006** |
| Corolla width (mm) | 25.51 ± 2.92 | 11.84 | 34.89 | 0.34 | 44.43 | < **0.001** |
| Flowering time | 44.87 ± 2.92 | 25.80 | 71.61 | 0.36 | 51.06 | < **0.001** |
| **Week 9** |  |  |  |  |  |  |
| Stolon number | 7.60 ± 0.76 | 0.80 | 3.69 | 0.22 | 21.02 | < **0.001** |
| Stolon length (mm) | 66.72 ± 10.4 | 319.96 | 895.46 | 0.36 | 55.06 | < **0.001** |
| Stem thickness (mm) | 2.45 ± 0.27 | 0.13 | 0.55 | 0.24 | 23.71 | < **0.001** |
| Flower number | 9.05 ± 2.21 | 23.06 | 51.21 | 0.45 | 91.19 | < **0.001** |
| Branch number | 6.24 ± 0.82 | 1.52 | 4.98 | 0.30 | 37.78 | < **0.001** |
| Node number | 11.25 ± 0.86 | 2.81 | 6.77 | 0.42 | 72.64 | < **0.001** |

**Table 2.** Loadings from the Principal Component Analysis (PCA) of phenotypic traits measured in the base generation. Trait loadings are shown for the first three principal components, with the proportion of total phenotypic variance explained by each component indicated in parentheses. Greater absolute values indicate a stronger contribution of the trait to that principal component.

| Trait | PC1 (28.7%) | PC2 (15.8%) | PC3 (9.1%) |
| --- | --- | --- | --- |
| **Week 5** |  |  |  |
| Early stolon number | 0.35 | 0.19 | 0.25 |
| Early stolon length | 0.29 | 0.33 | 0.18 |
| Corolla length | 0.32 | -0.04 | -0.60 |
| Corolla width | 0.32 | -0.04 | -0.59 |
| Flowering time | -0.27 | 0.38 | -0.29 |
| **Week 9** |  |  |  |
| Late stolon number | 0.09 | 0.44 | 0.09 |
| Late stolon length | 0.17 | 0.46 | 0.09 |
| Stem thickness | 0.31 | 0.33 | -0.03 |
| Flower number | 0.40 | -0.35 | 0.11 |
| Branch number | 0.20 | -0.21 | 0.23 |
| Node number | 0.43 | -0.18 | 0.14 |

### Univariate Response to Selection

Stolon number showed a clear response to selection after three rounds of artificial selection (generation x selection line: ^2^ = 67.78, df = 12, *P* < 0.001; Figure 2A; Table 3). Replicate selection lines within each direction (i.e., **H1** and **H2**, **L1** and **L2**) did not differ significantly across generations. In the high selection lines, plants produced 3.79 ± 0.09 (**H1**) and 3.86 ± 0.08 (**H2**) stolons before selection, increasing to 4.08 ± 0.13 (**H1**) and 4.39 ± 0.12 (**H2**) by generation four. Conversely, the low lines produced 3.54 ± 0.08 (**L1**) and 3.51 ± 0.09 (**L2**) stolons before selection and 3.17 ± 0.11 (**L1**) and 3.57 ± 0.12 (**L2**) stolons in generation four (Figure 2A; Table 3). The divergence between the groups was more pronounced when measured at later developmental stages (week 9), with high lines producing 5.83 ± 0.14 (**H1**) and 6.19 ± 0.13 (**H2**) stolons, compared 4.59 ± 0.13 (**L1**) and 4.79 ± 0.14 (**L2**) stolons in the low lines (Figure 2B; Table 4).

**Figure 2.**
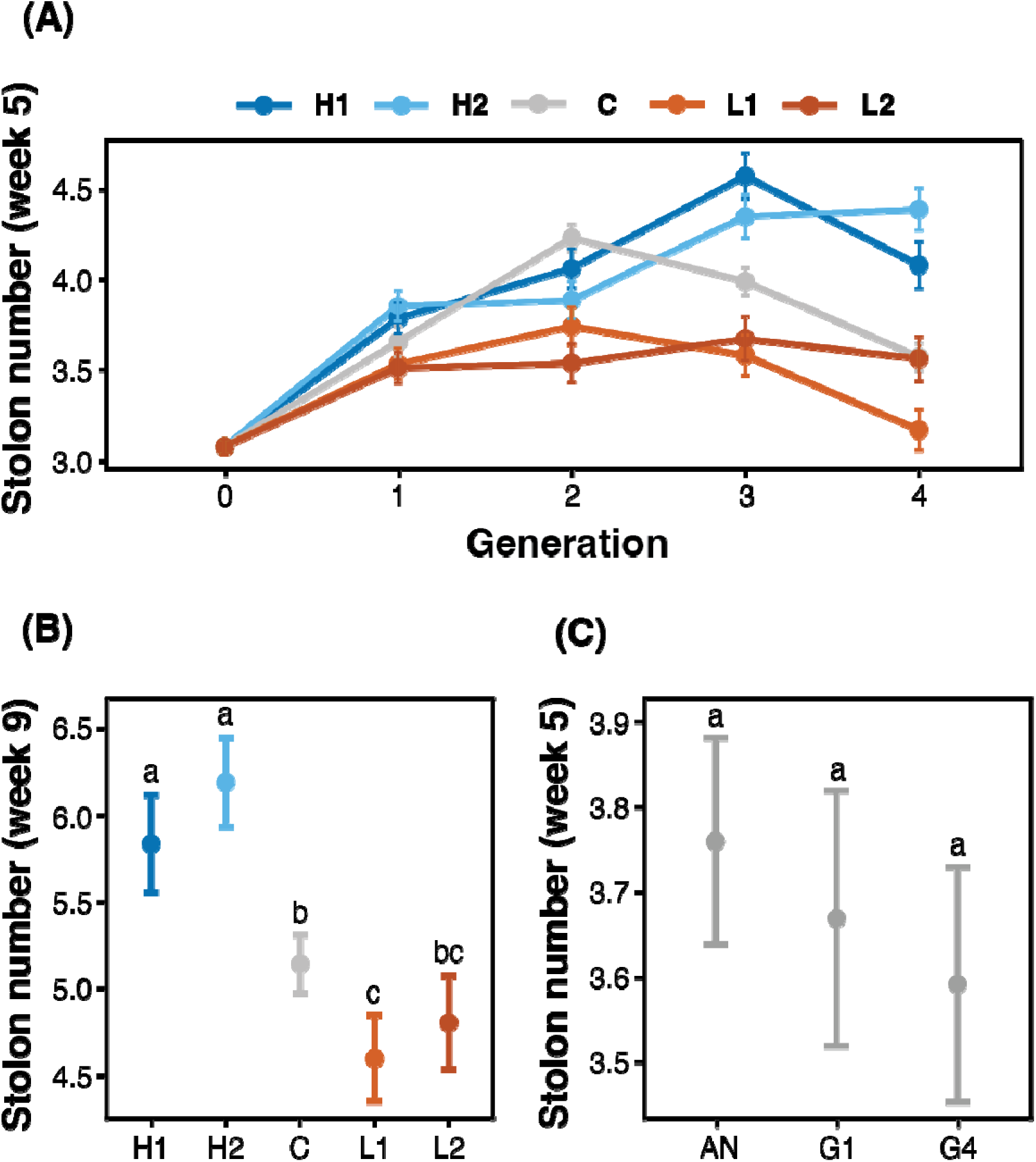
Change in stolon number across generations and selection lines. (A) Estimated marginal means (EMMs) ± 95% confidence intervals for stolon number at week 5 for each selection line (**H1**, **H2** = high; **C** = control; **L1**, **L2** = low) across four generations of selection. (B) EMMs ± 95% confidence intervals for late (week 9) stolon number in generation 4 plants across selection lines. (C) EMMs ± 95% confidence intervals for stolon number at week 5 in the ancestral population resurrected in generation 4 (**AN**), the entire first generation before selection (**G1**), and the control line in generation 4 (**G4**). Letters above points indicate groups that are significantly different from one another based on Tukey-adjusted pairwise comparisons (*P* < 0.05).

**Table 3.** Effects of artificial selection on stolon number in *Mimulus guttatus*, testing differences between control, high and low selection lines within each generation. A total of 10 pairwise contrasts were performed within each generation, with an additional five contrasts in the final generation. *P*-values were adjusted for multiple comparisons using the Tukey’s method correction; bold values indicate statistically significant differences.

| Pairwise contrasts | Generation 1 |  |  |  | Generation 2 |  |  |  | Generation 3 |  |  |  | Generation 4 |  |  |  |
| --- | --- | --- | --- | --- | --- | --- | --- | --- | --- | --- | --- | --- | --- | --- | --- | --- |
|  | Est | SE | t-stat | *P*-value | Est | SE | t-stat | *P*-value | Est | SE | t-stat | *P*-value | Est | SE | t-stat | *P*-value |
| H1 vs. H2 | -0.668 | 0.11 | -6.03 | 0.97 | 0.175 | 0.15 | 1.18 | 0.77 | 0.221 | 0.17 | 1.3 | 0.69 | -0.307 | 0.17 | -1.78 | 0.48 |
| L1 vs. L2 | 0.025 | 0.11 | 0.2 | 0.99 | 0.203 | 0.14 | 1.38 | 0.64 | -0.69 | 0.16 | -4.55 | 0.98 | -0.393 | 0.16 | -2.36 | 0.17 |
| H1 vs. L1 | 0.252 | 0.11 | 2.31 | 0.14 | 0.32 | 0.14 | 2.17 | 0.19 | 0.988 | 0.17 | 5.9 | **<0.001** | 0.916 | 0.17 | 5.35 | **<0.001** |
| H1 vs. L2 | 0.273 | 0.1 | 2.51 | 0.09 | 0.524 | 0.15 | 3.51 | **0.0041** | 0.898 | 0.17 | 5.19 | **<0.001** | 0.517 | 0.18 | 2.92 | **0.04** |
| H2 vs. L1 | 0.321 | 0.11 | 2.95 | 0.03 | 0.145 | 0.14 | 0.99 | 0.86 | 0.766 | 0.16 | 4.77 | **<0.001** | 1.216 | 0.16 | 7.5 | **<0.001** |
| H2 vs. L2 | 0.342 | 0.11 | 3.14 | 0.01 | 0.348 | 0.15 | 2.34 | 0.13 | 0.677 | 0.17 | 4 | **0.0005** | 0.824 | 0.16 | 4.86 | **<0.001** |
| C vs. H1 | -0.128 | 0.11 | -1.19 | 0.76 | 0.168 | 0.13 | 1.3 | 0.69 | -0.579 | 0.15 | -3.99 | **0.0006** | -0.508 | 0.15 | -3.42 | **0.01** |
| C vs. H2 | -0.198 | 0.1 | -1.82 | 0.36 | 0.343 | 0.13 | 2.66 | 0.06 | -0.358 | 0.14 | -2.55 | 0.08 | -0.815 | 0.14 | -5.84 | **<0.001** |
| C vs. L1 | 0.122 | 0.1 | 1.15 | 0.79 | 0.488 | 0.12 | 3.83 | **0.001** | 0.408 | 0.13 | 3.07 | 0.02 | 0.402 | 0.14 | 2.96 | **0.04** |
| C vs. L2 | 0.144 | 0.11 | 1.2 | 0.99 | 0.691 | 0.13 | 5.83 | **<0.001** | 0.319 | 0.14 | 2.25 | 0.17 | 0.009 | 0.14 | 0.06 | 1 |
| Ancestor vs. H1 | - | - | - | - | - | - | - | - | - | - | - | - | -0.541 | 0.14 | -2.56 | 0.17 |
| Ancestor vs. H2 | - | - | - | - | - | - | - | - | - | - | - | - | -0.648 | 0.13 | -4.8 | **<0.001** |
| Ancestor vs. C | - | - | - | - | - | - | - | - | - | - | - | - | 0.167 | 0.1 | 1.64 | 0.57 |
| Ancestor vs. L1 | - | - | - | - | - | - | - | - | - | - | - | - | 0.569 | 0.13 | 4.344 | **0.0002** |
| Ancestor vs. L2 | - | - | - | - | - | - | - | - | - | - | - | - | 0.176 | 0.14 | 1.26 | 0.821 |

**Table 4.** Pairwise contrasts of Estimated Marginal Means (EMMs) for stolon number at week 9 (corresponding to Figure 2B) in generation 4. A total of 10 pairwise comparisons were performed among selection lines (control, high and low). *P*-values were adjusted for multiple comparisons using Tukey’s method, and significant differences (*P* < 0.05) are shown in bold.

| Pairwise contrasts | Estimate | SE | *t*-statistic | *P*-value |
| --- | --- | --- | --- | --- |
| C vs. H1 | -0.695 | 0.166 | -4.192 | **0.0003** |
| C vs. H2 | -1.050 | 0.156 | -6.736 | < **0.0001** |
| C vs. L1 | 0.543 | 0.152 | 3.585 | **0.0035** |
| C vs. L2 | 0.338 | 0.161 | 2.102 | 0.2219 |
| H1 vs. H2 | -0.355 | 0.193 | -1.839 | 0.3528 |
| H1 vs. L1 | 1.239 | 0.190 | 6.522 | < **0.0001** |
| H1 vs. L2 | 1.034 | 0.198 | 5.233 | < **0.0001** |
| H2 vs. L1 | 1.593 | 0.181 | 8.789 | < **0.0001** |
| H2 vs. L2 | 1.388 | 0.189 | 7.336 | < **0.0001** |
| L1 vs. L2 | -0.205 | 0.186 | -1.105 | 0.8038 |

The selection lines tended to deviate from the control line in the expected directions and were significantly different from the opposing selection lines, though two exceptions exist. First, the high line plants produced slightly fewer stolons than the control line in generation 2, with the high lines producing 4.06 ± 0.11 (**H1**) and 3.89 ± 0.11 (**H2**) stolons, and the control line individuals producing 4.23 ± 0.07 stolons (Figure 2A; Table 3). Second, in generation 4, the low lines did not produce significantly fewer stolons compared to the control (Figure 2A; Table 3).

Realized heritabilities for the high selection lines were generally larger than those for the low lines, although they were quite variable for all groups (Table 5). Both the second high and second low lines showed very little response to selection in generation 1 and generation 3, and subsequently had realized heritabilities near zero; however, both showed an increase in selection response and heritability in generation 2 and generation 4 (Table 5).

**Table 5.** Realized ***h^2^*** ± SE for each of three generations of selection on stolon number in *M. guttatus*.

| Generation | High 1 | High 2 | Low 1 | Low 2 |
| --- | --- | --- | --- | --- |
| 1 | 0.353 ± 0.178 | 0.063 ± 0.187 | -0.243 ± 0.179 | -0.028 ± 0.193 |
| 2 | 0.584 ± 0.175 | 0.491 ± 0.166 | 0.154 ± 0.155 | -0.131 ± 0.158 |
| 3 | -0.447 ± 0.186 | 0.057 ± 0.182 | 0.377 ± 0.150 | 0.094 ± 0.177 |

To test for inadvertent selection and/or genetic drift over the course of the experiment, we compared three groups: the control group which underwent four generations in the greenhouse with random mating each generation; a resurrected ancestral group grown alongside the fourth-generation plants; and the entire first generation of plants before selection. Across these groups, we found little indication that there was inadvertent selection or drift on stolon number: the mean number of stolons at week five did not differ significantly (F_2,1295_ = 2.24, *P* = 0.107; Figure 3C; Table S5) between the entire first generation (estimated marginal mean ± SE = 3.67 ± 0.08; N = 1678), the ancestral population grown in generation four (3.76 ± 0.06; N = 509), and the fourth-generation control line (3.59 ± 0.07; N = 515).

**Figure 3.**
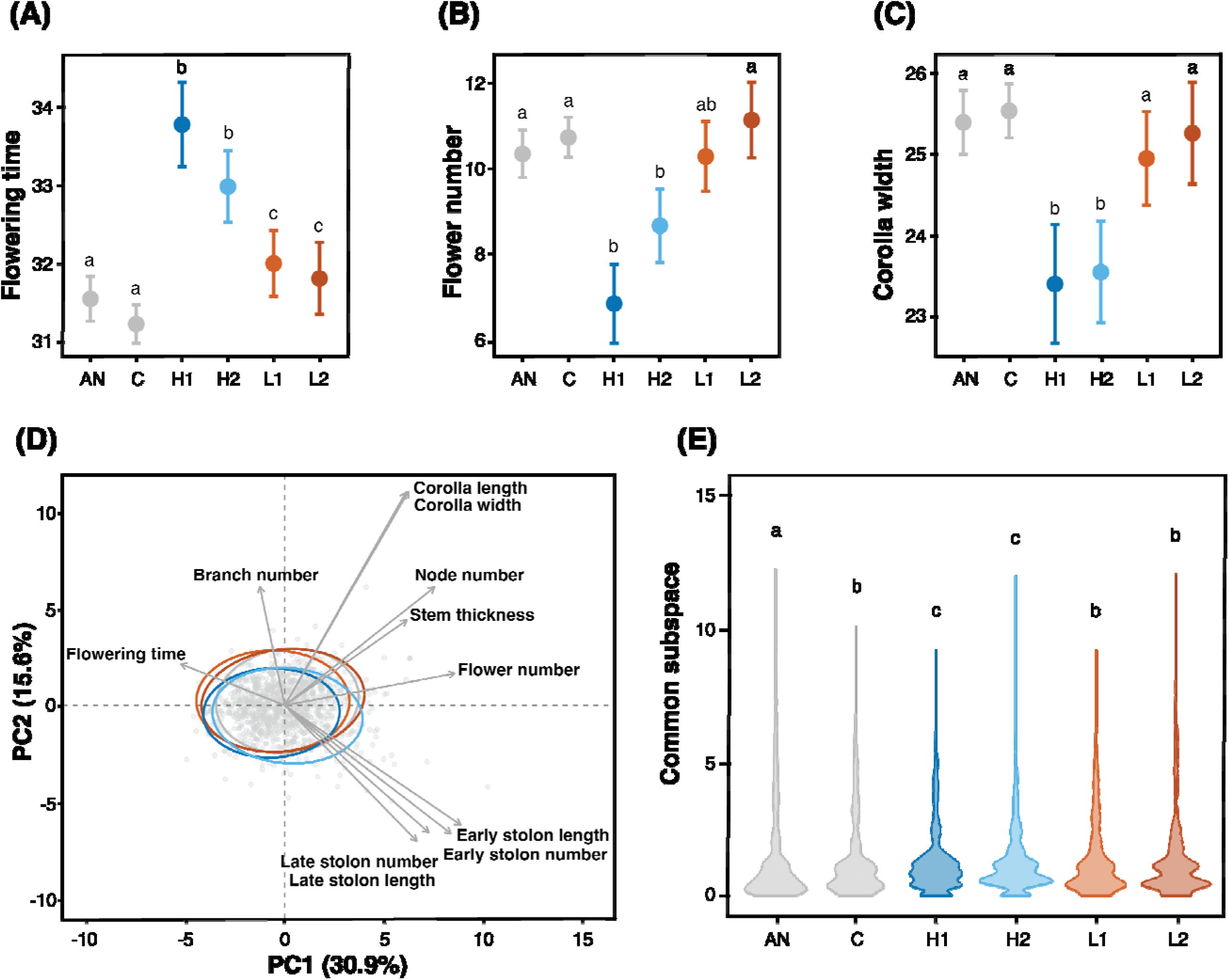
Multivariate response to artificial selection. (A-C) Estimated Marginal Means (EMMs) ± 95% confidence intervals of floral traits (A: flowering time, B: flower number, C: corolla width) in generation 4 across selection lines, including the ancestral population. Points represent model-adjusted group means with error bars indicating 95% confidence intervals. Letters above points indicate groups that are significantly different from one another based on Tukey-adjusted pairwise comparisons (*P* < 0.05). (D) Principal Component Analysis (PCA) of phenotypic trait variation in generation 4 from five selection lines (**H1**, **H2**, **C**, **L1**, **L2**). Each point represents an individual, with 95% confidence ellipses shown for each group. Arrows indicate the loadings of traits on PC1 and PC2 (E) Distributions of Krzanowski’s common subspace values for each selection line, including the ancestral population (**AN**), grown in generation 4. Violin plots display the distribution of common subspace values from 1000 resampled ***G*** matrices per group. Larger values indicate greater similarity in the leading eigenvectors of the ***G*** matrices, representing conserved directions of genetic variation.

### Multivariate Response to Selection

Phenotypic analyses of multivariate patterns in generation 4 using linear-mixed effects models revealed that high selection line plants flower substantially later (Figure 3A), have fewer flowers (Figure 3B), and produce smaller flowers (Figure 3C) than both the control and low lines. The PCA revealed multivariate phenotypic structure (Figure 3D; Table 6). Once again, PC1, which explained 30.9% of the total phenotypic variance, captured variation in overall plant size, with both stolon traits and flowering time having positive loadings (Figure 3D; Table 6). A trade-off between clonality and flowering was described by PC2 (15.6% of variance) with high positive loadings for floral traits, and negative loadings for clonal traits (Figure 3D; Table 6); and both the control and low lines had greater values on PC2 than high lines (F_4,1080_ = 19.52, *P* < 0.001; Table S6).

**Table 6.**
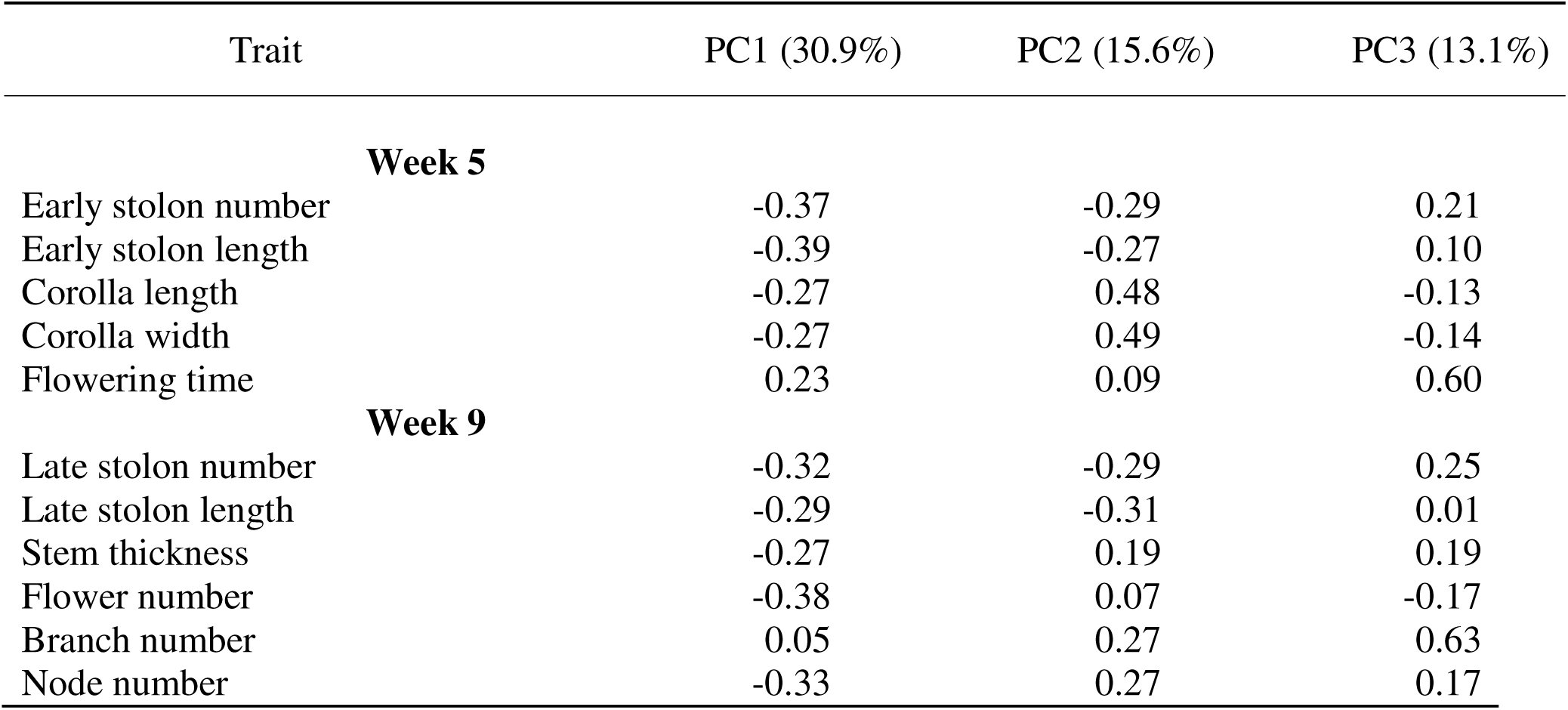
Loadings from the Principal Component Analysis (PCA) of standardized phenotypic traits measured across selection lines in generation 4. Trait loadings are shown for the first three principal components, with the proportion of total phenotypic variance explained by each component indicated in parentheses. Greater absolute values indicate a stronger contribution of the trait to that principal component.

| Trait | PC1 (30.9%) | PC2 (15.6%) | PC3 (13.1%) |
| --- | --- | --- | --- |
| **Week 5** |  |  |  |
| Early stolon number | -0.37 | -0.29 | 0.21 |
| Early stolon length | -0.39 | -0.27 | 0.10 |
| Corolla length | -0.27 | 0.48 | -0.13 |
| Corolla width | -0.27 | 0.49 | -0.14 |
| Flowering time | 0.23 | 0.09 | 0.60 |
| **Week 9** |  |  |  |
| Late stolon number | -0.32 | -0.29 | 0.25 |
| Late stolon length | -0.29 | -0.31 | 0.01 |
| Stem thickness | -0.27 | 0.19 | 0.19 |
| Flower number | -0.38 | 0.07 | -0.17 |
| Branch number | 0.05 | 0.27 | 0.63 |
| Node number | -0.33 | 0.27 | 0.17 |

Permutation tests comparing ***G*** matrix sizes among groups revealed significant differences (F_5,5994_ = 158.8, *P* < 0.001; Figure S4; Table S7). With the exception of **L2**, all groups had smaller ***G*** matrices than the ancestral population. Furthermore, the selected groups (**H1**, **H2**, **L1**, **L2**) exhibited larger ***G*** matrix sizes than the control line, suggesting that genetic drift may have occurred in the control line over the course of the experiment, or selecting on a single trait expanded the genetic variation on other traits (Figure S4; Table S7).

When examining ***G*** matrix subspaces, we found significant divergence from the ancestral population by all groups, suggesting that multivariate phenotypes changed over time for all selection lines (Figure 3E; Table S8). Furthermore, relative to the control, significant divergence was detected only in the high selection lines. No significant differences were observed between replicates within selection directions, but there was a significant difference when comparing high and low selection lines (Figure 3E; Table S8). Overall, artificial selection for increased stolon number appeared to have the strongest effect on ***G*** matrix variance, with the high selection lines showing the most pronounced divergence both phenotypically (Figure 3A) and in genetic variance-covariance space (Figure 3E; Table S8).

## DISCUSSION

Here we show extensive, heritable variation in stolon production within a single population of *Mimulus guttatus* grown in a common greenhouse environment and demonstrate that artificial selection produces a clear response. We further demonstrate genetic and phenotypic correlations between clonal traits and sexual traits including flowering time and floral display. After four generations of divergent selection for increased or decreased stolon number, we show that a single population of *M*. *guttatus* has the genetic capacity to respond and evolve; plants in the high lines produced significantly more stolons than plants in the control and low lines. Similarly, plants from the high and low selection lines differed significantly in traits correlated with stolon number: high line plants flowered later, produced fewer flowers, and had smaller flowers than both the low line, control line, and ancestral population. In contrast, plants from the low selection line showed a more muted response and remained largely similar to the control and ancestral population in both clonal and sexual trait means. This difference is further highlighted by patterns of genetic covariation, with high lines showing much greater changes in both the size and shape of their ***G*** matrices compared to low lines, suggesting that selection for increased clonality could lead to a change in the multivariate phenotype in *M. guttatus*. These results demonstrate that artificial selection on clonality can rapidly reshape trait distributions and multivariate genetic architecture and can enhance or diminish genetic correlations that may affect evolutionary trajectories in natural populations.

### Realized Heritability and Response to Selection

Studies on quantitative genetic variation and heritability of clonal life-history traits are scarce (Fischer and van Kleunen, 2002). Indeed, most studies on the life history of *M. guttatus* have focussed on sexual traits and demonstrated heritable variation in flower size (van Kleunen and Ritland, 2005; Monnahan and Kelly, 2017), flowering time (Robertson et al., 1994; Monnahan and Kelly, 2017), sex allocation (Ritland and Ritland, 1989), and mating system (van Kleunen and Ritland, 2004; van Kleunen and Ritland, 2005). Compared to these sexual traits, the heritability of vegetative traits, including stolon number tends to be lower. A previous estimate of broad-sense heritability for stolon number in an outbred greenhouse population is 0.31 (Rubin et al., 2018), which is comparable to our estimate in the base population of 0.25 (although an estimate of 0.71 was found in a hybrid F2 population: Coughlan et al., 2021), and similar to estimates for stolons in *Trifolium repens* (Caradus and Chapman, 1996). One explanation for the lower heritability is that stolon number is determined by multiple quantitative trait loci that are further influenced by genotype-by-environment interactions. Unlike flowering time which typically involves only a few major-effect loci (Hall et al., 2006; Hall and Willis, 2006; Kelly and Mojica, 2011), investment in stolons involves a meristematic trade off (Baker et al., 2012) and additional allocation to vegetation (e.g., internode elongation, nodes, stolon leaves).

Despite the moderate heritability in the base population, we elicited a rapid, but asymmetrical change in the number of stolons with artificial selection. Low lines diverged from control less than high lines and had more variable and lower realized heritability. This asymmetrical response was not necessarily expected, but several processes may account for it. First, individuals that make few stolons could do so because they carry particular allelic variants for the trait, or they may make few stolons because they are overall lower in quality and less vigorous. Plants may be lower quality for environmental reasons related to small-scale variation in the greenhouse (i.e., minor differences in light or water availability); or for genetic reasons that are unrelated to stolon investment (i.e., differences in overall condition). If any of these processes are an underlying cause of reduced stolon investment, then it would undermine our ability to select on the trait itself, resulting in lower heritability and response. Importantly, this cause of the difference in stolon investment is likely to be found in nature as well and may contribute to differences in evolutionary responses to natural selection.

Another potential explanation for a weaker response in the low lines is that four generations of selection were not enough to elicit an evolutionary response in this direction. In an experiment involving 10 generations of selection on anther exsertion in *Raphanus raphanistrum,* low lines did not diverge from control plants, and sometimes had greater mean trait values during intermediate generations (Conner et al., 2011). Similarly, other artificial selection experiments have failed to show a response to selection in the first three or four generations (Semlitsch and Wilbur, 1989; van Kleunen et al., 2002; Fischer et al., 2003; Tisinai and Busch, 2024). Finally, the degree of response to selection could be impeded by inadvertent selection in the greenhouse, inbreeding depression, and/or genetic drift. While all of these processes should affect the low, high, and control, lines equally, it is possible that the phenotypic effects manifest as fewer stolons. Nonetheless, at the end of the experiment when we grew the control line and ancestral populations simultaneously, we found that they did not differ and had similar distributions of stolon number. This suggests that inbreeding and/or genetic drift were unlikely to have had a significant effect on stolon number in this experiment.

### Multivariate Response to Selection

Patterns of covariation were fairly similar in all lines after selection, as we observed an almost complete overlap in both trait means and ellipses along PC1 and PC2 between the control and low lines; however, the high lines diverged almost exclusively due to stolon-related traits. Similarly, high line plants diverged from control and low lines in their ***G*** matrix structure, as evidenced by the Krzanowski’s common subspace analysis. Finally, high line plants maintained stronger correlations between stolon number and floral traits than either low lines or the control, reflecting a greater magnitude of phenotypic and genetic (co)variation in the high lines. Previous selection experiments looking at sexual dimorphism and floral display in *Silene* (Delph et al., 2007) and bract size and flower size in *Dalechampia* (Hansen et al., 2003) revealed that even in the presence of substantial additive genetic variation for traits under selection, genetic correlations among traits can constrain their evolvability, thus limiting the response to unidirectional selection. Taken together, the relative response in stolon number and multivariate change in the high lines relative to the low lines suggest that selection against clonality was constrained by existing covariation (so that the low lines remained similar to the control), whereas selection for increased clonality drove divergence along a major axis of trait variation dominated by clonality.

These multivariate patterns underscore how genetic covariation can channel evolutionary trajectories. The divergence of high lines along an axis dominated by clonality, contrasted with the lack of change in low lines, illustrates how responses depend on the orientation of standing variation. Similar outcomes have been reported in both selection experiments and natural populations, where ***G*** matrix structure tends to evolve slowly and respond to selection along major axes of genetic variance rather than in the exact direction of selection (Arnold et al., 2008; Lind et al., 2015). At the same time, a variety of studies have also shown that selection can reshape ***G*** even over short timescales (Fong, 1989; Roff et al., 2004; Steven et al., 2007; Doroszuk et al., 2008), highlighting the dynamic nature of genetic constraints. Our results are consistent with both views, showing that selection on traits harbouring substantial additive genetic variation, particularly those aligned with major axes of genetic variance, may produce more rapid changes in ***G***, whereas traits with less variation may evolve more slowly.

Together, this suggests that clonality is not an independently evolving trait, but part of a broader multivariate suite linking vegetative and reproductive traits. This emphasizes that the predictability of evolutionary response depends not only on the amount of genetic variation but also on the structure and stability of the ***G*** matrix across generations. Future work estimating ***G*** in natural populations of *M. guttatus* will be essential for understanding how eigenstructures are maintained or reshaped under multiple selective pressures, and how this constrains or facilitates the evolution of clonality.

While selection altered mean trait values for stolons in the expected directions, it also reshaped the structure of trait correlations. In the base generation, stolon number was strongly correlated with flowering time, but after selection this correlation weakened, particularly in the low lines. This suggests that selection can quickly erode genetic correlations, either through multiple rounds of recombination reducing linkage, or by selecting against pleiotropic loci, or through the evolution of new correlations. For example, selection on flower size in the sexually dimorphic plant *Silene latifolia* not only reduced correlated responses in flowers of either sex but also uncovered new correlations between flower size and number (Delph et al., 2007). Similar results have been found in other species: artificial selection on flowering time in *Arabidopsis thaliana* altered correlations with leaf number and rosette size (Méndez-Vigo et al., 2012), and domestication of maize reshaped correlations between reproductive and vegetative traits as selection on ear and kernel traits reduced associations with plant architecture (Yang et al., 2019). The high lines in this experiment maintained stronger correlations between stolon number and flower number than low lines, potentially due to the greater phenotypic and genetic variation within the high lines, or due to the alignment between the direction of selection and the major axes of genetic covariation.

### Implications for Perennial Evolution

Evolutionary biologists have long used domestication to study the implications of selection under controlled conditions (Darwin, 1899; Ross-Ibarra et al., 2007; Pickersgill., 2009), and in the case of herbaceous plants this has largely involved the breeding of annual species. Although recent research has focussed on developing perennial crops (Cox et al., 2006; DeHaan et al., 2023), we still know comparatively little about the fundamental genetic, physiological, and developmental differences between herbaceous annual and perennial species (Lundgren and Des Marais, 2020). The diversity of life-history strategies within *M. guttatus* – ranging from diminutive annuals to large, clonal perennials, provides the raw genetic material to understand the evolution of traits that comprise these life-history strategies. Among our initial motivations was to determine if we could select on a single population to recreate both annual-like phenotypes (low stolon number) and/or more clonal phenotypes (high stolon number). While three generations of artificial selection were not sufficient to produce these extremes, we did find that the selection lines had a series of correlated traits consistent with those found in nature.

In studying trait covariation in annual and perennial herbaceous crops, researchers have found that the developmental stage of the individual can affect patterns of variation. In particular, when examining mature plants, correlations between vegetative and reproductive traits were overwhelmingly positive and perennial species exhibited more numerous positive correlations between traits than annual species (Rubin et al., 2024); while studies of juvenile crop species found correlations between vegetative and reproductive traits were stronger for annuals than perennials (Herron et al., 2021). In line with this, we found different patterns of correlated traits at different developmental stages. Prior to reproduction (i.e., at week 5), plants tended to display positive correlations amongst all vegetative traits and stolons. As plants continued to mature and enter the reproductive stage, they expressed more variation in their trait values and new correlations emerged. For example, pre-flowering (i.e., at week five) plants with many stolons were typically larger than plants with few stolons, whereas reproductive adults (i.e., at week nine) with many stolons were outgrown by these plants and were typically smaller than plants with few stolons. Although trait correlations shifted between week five and week nine, these changes are unlikely to meaningfully bias selection outcomes, as early differences largely reflected overall growth trajectories rather than shifts in trait rankings. In line with this, plants producing many stolons were also those that flowered later, a result that has been previously described in this species (Hall et al., 2006; Friedman et al., 2015; Rubin et al., 2018).

One of the underlying components of perenniality is the maintenance of at least one meristem beyond the first growing season, to facilitate growth and reproduction in subsequent years (Friedman, 2020). One mechanism to maintain meristem indeterminacy is clonality, as seen in *M. guttatus*, where meristems are spread out through space and time by the growth of genetically identical stems. The genetic basis of these perennating structures may involve only a few QTL (Friedman et al., 2015) and include genes controlling growth hormones involving gibberellin (Olsen et al., 2025; Kollar et al., 2025). Gibberellin is a phytohormone responsible for flower development and the elongation and expansion of vegetative traits (Blasquez et al., 1998; Richards et al., 2001) and has been shown in *M. guttatus* to affect bolting and the proliferation of stolons (Lowry et al., 2019; Olsen et al., 2025). Indeed, hormonal regulation of flowering versus vegetative growth may be a more general feature of perennial herbaceous plants. Both *Lolium* and *Chrysanthemum* show similar patterns: plants that are dosed with gibberellin in early life stages bolt and flower much earlier and invest less in vegetation in later life (King et al., 2003; Yang et al., 2014). While it is beyond the scope of this study to identify the genetic or physiological causes of the differences in phenotypes, it is possible that the artificial selection we performed here altered allele frequencies at genes related to hormone regulation and/or genes controlling meristem fate.

## Conclusion

The observed change in traits and trait correlations over a microevolutionary time scale has consequences for our understanding of macroevolutionary patterns. Testing whether these dynamics increase or decrease in natural settings is essential for evaluating their contribution to broad scale evolutionary patterns. More research is needed to determine the extent to which our results apply to wild populations experiencing spatial-temporal variation in natural selection gradients, as well as natural rates of non-random mating, immigration and emigration, and fluctuations in population size. Assuming some level of similarity in greenhouse and wild population ***G*** matrix evolutionary dynamics, our results have important implications for populations facing consistent and directional environmental change. Clonal reproduction in *Mimulus guttatus*, as well as in plants broadly, is a quantitative trait that has the capacity to affect the resilience and adaptive capacity of a population as it strongly affects the ability of individuals to reproduce, disperse, and persist over time. Stolon production and thus perenniality also results in strong local adaptation but weak foreign adaptation (in annual-like habitats) in *M. guttatus*, suggesting that directional selection can both strongly favour clonality and strongly select against it depending on habitat permanence (Lowry and Willis, 2010; DeMarche et al., 2017). This work demonstrates how populations harbouring extensive genetic variation are more likely to be resilient in the face of climate change and are capable of rapid adaptive evolution.

## Author Contributions

C.S. and J.F. conceived the project; C.S., I.L., J.L. collected the data; C.S. developed the code and analyzed the data; C.S. and J.F. wrote the manuscript; J.F. oversaw all aspects of the project. All authors approved the final version of the manuscript.

## Acknowledgements

The authors thank colleagues at Queen’s University, including W. van Drunen for guidance in developing code and collecting data; C. Smith and A. Van Natto for assistance with pollinations and interpretations of results; E. Gillette, and B. Balcaran for help collecting data; and the Queen’s University Phytotron team for maintaining the greenhouse during this project. The authors also thank colleagues C. Eckert, P. McKenzie, J. Stinchcombe, F. Wu for guidance on analyses.

## Data availability statement

Data and code will be available from Dryad DOI: 10.5061/dryad.

