## Supplemental information for "Life-history evolution under artificial selection in a clonal plant"

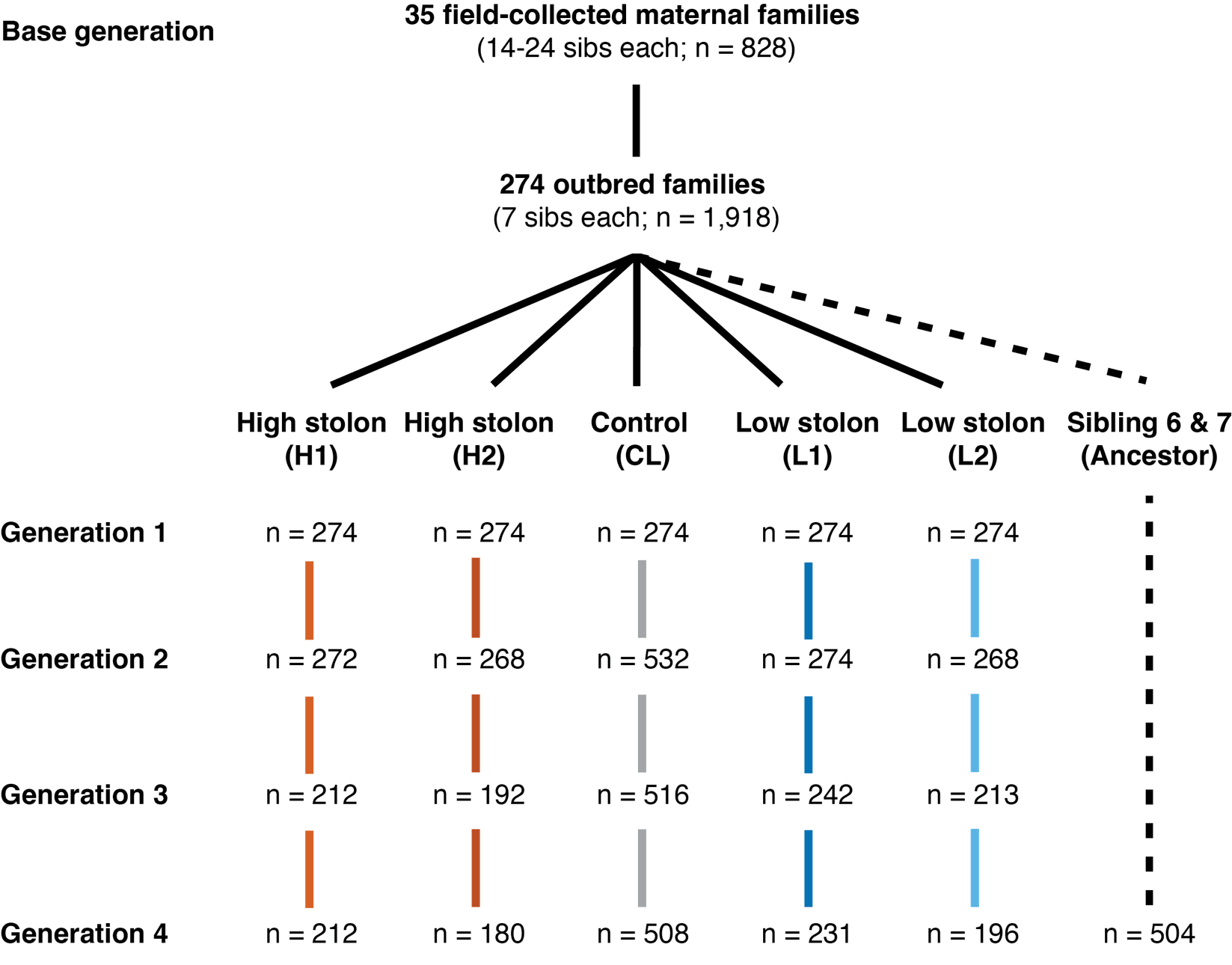

**Figure S1.** Artificial selection experimental design using *Mimulus guttatus*. Seed collected from the field was grown to create the base generation (**G0**), comprised of 35 open-pollinated maternal families (14-24 siblings each: n = 828). These plants were systematically paired and crossed to maximize different between-family crosses and maintain family representation, producing 274 outbred families. Full siblings from each outbred family were equally distributed into 6 groups: two replicate high lines, two replicate low lines, one control (with twice as many individuals after selection), and a stored ancestral line. Each high and low line underwent three rounds of selection on stolon number and length. Numbers along horizontal lines represent population sizes across selection lines within each generation.

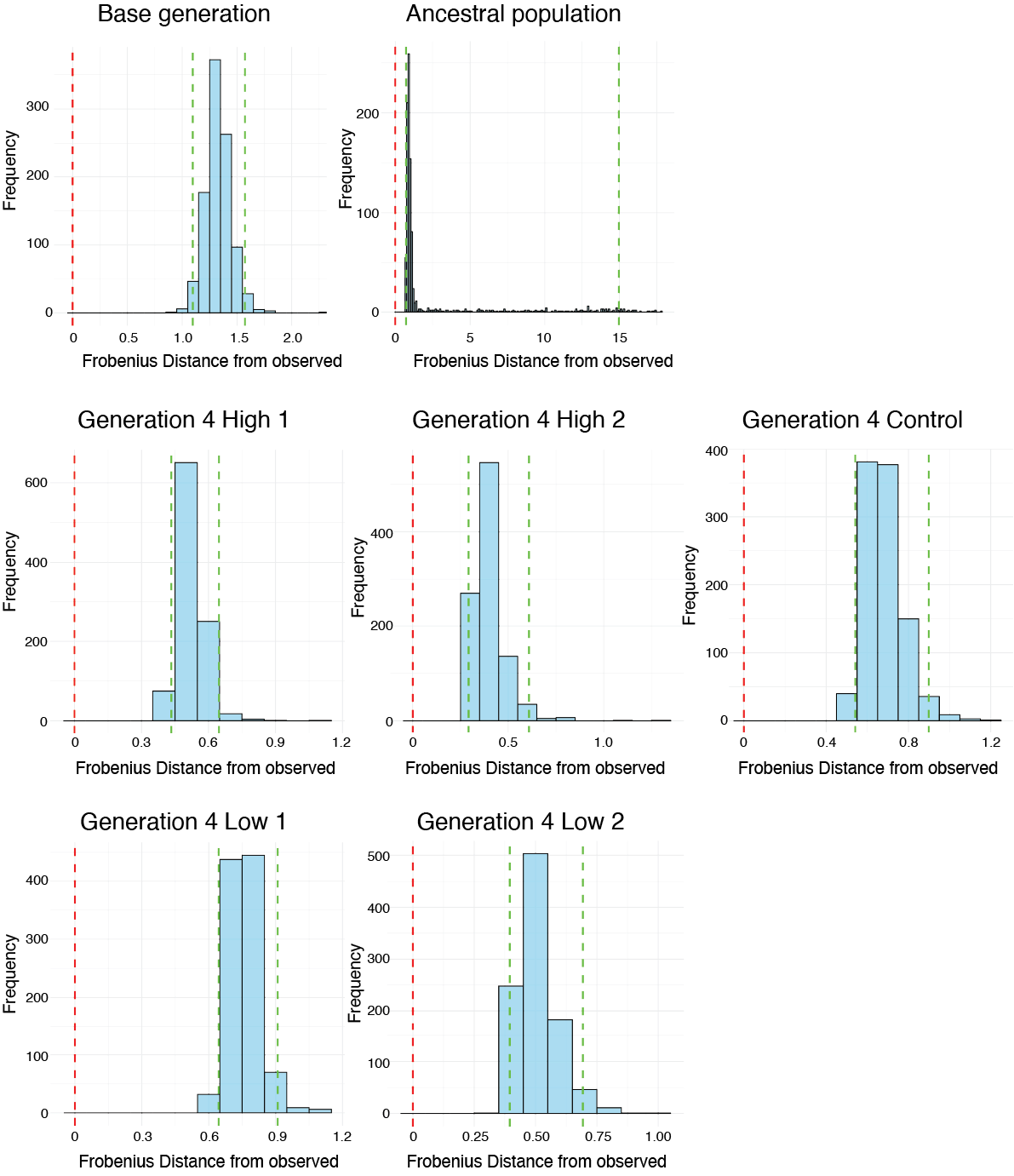

**Figure S2.** Null distributions of Frobenius distances comparing observed and randomized ***G*** matrices in the base and selection populations of *Mimulus guttatus*. Dashed green lines indicate the 95% quantile interval of the null distribution, and the dashed red line marks the Frobenius distance between the observed ***G*** matrix and itself (0). *P*-values were calculated as the proportion of null distances less than or equal to the observed ***G*** matrix from random expectation.

**Table S1**. Observed ***G*** matrix Frobenius distances from the observed ***G*** matrices, and the 95% confidence intervals of the randomized distributions of ***G*** matrices. Bold confidence intervals indicate the significance of the observed ***G*** matrix relative to its null distribution (*P* <0.001).

| Group |  | Frobenius distance from  the observed ***G*** matrix | | |
| --- | --- | --- | --- | --- |
|  |  | Observed difference | | 95% CI |
| Base generation (G0) |  | | 0 | **[1.10, 1.57]** |
| Ancestral population |  | | 0 | **[0.72, 14.95]** |
| Control |  | | 0 | **[0.54, 0.90]** |
| High 1 |  | | 0 | **[0.43, 0.65]** |
| High 2 |  | | 0 | **[0.29, 0.61]** |
| Low 1 |  | | 0 | **[0.64, 0.91]** |
| Low 2 |  | | 0 | **[0.39, 0.69]** |

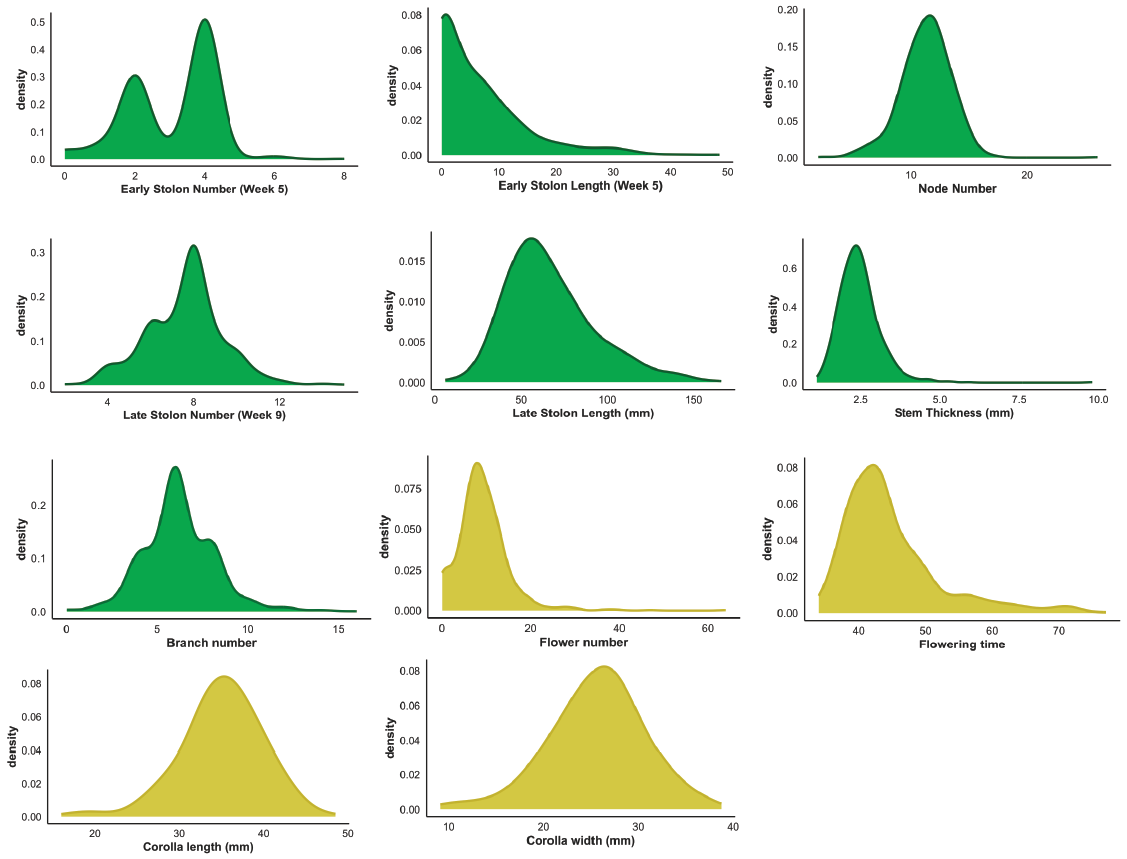

**Figure S3.** Density distributions of 11 phenotypic traits measured in the base generation for 828 *Mimulus guttatus* plants. Each panel shows the distribution of individual trait values across all plants in the base generation. Traits related to vegetative growth are shown in green, while traits related to reproductive characteristics are shown in yellow.

**Table S2.** Phenotypic correlation matrix (Pearson’s *ρ*) for 11 traits measured in the base generation (**G0**) on 828 *Mimulus guttatus* plants. Phenotypic correlations were calculated using fitted values from linear mixed models for each trait, with seed family included as a random effect and block as a fixed effect. The fitted values were extracted from the models and Pearson correlation coefficients were calculated on the fitted values. *P*-values for each correlation were generated using a bootstrapped significance test with a 95% confidence interval. Significant values are bolded (*P* < 0.05).

|  |  | **Early stolon number** | | | **Early stolon length** | | **Corolla length** | | **Corolla width** | | **Days to flower** | | **Late stolon  number** | | **Late stolon  length** | | **Stem thickness** | | **Flower number** | | **Branch number** | | **Node number** |
| --- | --- | --- | --- | --- | --- | --- | --- | --- | --- | --- | --- | --- | --- | --- | --- | --- | --- | --- | --- | --- | --- | --- | --- |
| **Early stolon number** | | | 1 | **0.632** | | **-0.071** | | 0.002 | | -0.057 | | **0.270** | | **0.248** | | **0.176** | | **-0.187** | | **0.224** | | **-0.164** | |
| **Early stolon length** | | | **0.632** | 1 | | 0.032 | | 0.026 | | **-0.143** | | **0.351** | | **0.430** | | **0.113** | | **-0.133** | | 0.046 | | **-0.113** | |
| **Corolla length** | | | **-0.071** | 0.032 | | 1 | | **0.871** | | **-0.362** | | **-0.225** | | **-0.106** | | 0.046 | | **0.490** | | **0.099** | | **0.330** | |
| **Corolla width** | | | 0.002 | 0.026 | | **0.871** | | 1 | | **-0.293** | | **-0.198** | | -0.043 | | **0.145** | | **0.417** | | **0.122** | | **0.276** | |
| **Days to flower** | | | -0.057 | **-0.143** | | **-0.362** | | **-0.293** | | 1 | | **0.384** | | **0.141** | | **0.417** | | **-0.736** | | **-0.311** | | **-0.521** | |
| **Late stolon number** | | | **0.270** | **0.351** | | **-0.225** | | **-0.198** | | **0.384** | | 1 | | **0.263** | | **0.403** | | **-0.412** | | **-0.335** | | **-0.167** | |
| **Late stolon length** | | | **0.248** | **0.430** | | **-0.106** | | -0.043 | | **0.141** | | **0.263** | | 1 | | **0.511** | | **-0.148** | | **-0.396** | | -0.020 | |
| **Stem thickness** | | | **0.176** | **0.113** | | 0.046 | | **0.145** | | **0.417** | | **0.403** | | **0.511** | | 1 | | **-0.126** | | **-0.412** | | 0.111 | |
| **Flower number** | | | **-0.187** | **-0.133** | | **0.490** | | **0.417** | | **-0.736** | | **-0.412** | | **-0.148** | | **-0.126** | | 1 | | **0.329** | | **0.781** | |
| **Branch number** | | | **0.224** | 0.046 | | **0.099** | | **0.122** | | **-0.311** | | **-0.335** | | **-0.396** | | **-0.412** | | **0.329** | | 1 | | **0.280** | |
| **Node number** | | | **-0.164** | **-0.113** | | **0.330** | | **0.276** | | **-0.521** | | **-0.167** | | -0.020 | | 0.111 | | **0.781** | | **0.280** | | 1 | |

**Table S3.** Genetic variance-covariance (***G***) matrix for 11 traits measured in the base generation (**G0**) on 828 *Mimulus guttatus* plants. values exceeding 1 on the diagonal elements indicate that the genetic variance for that specific trait is greater than the phenotypic variance, meaning there is a larger potential for evolutionary change in that trait due to genetic factors alone; while off-diagonal values over 1 suggest a strong positive genetic covariance between two traits, implying that selection on one trait could significantly influence the evolution of the other.

|  |  | **Early stolon number** | | | **Early stolon length** | | **Corolla length** | | **Corolla width** | | **Days to flower** | | **Late stolon  number** | | **Late stolon  length** | | **Stem thickness** | | **Flower number** | | **Branch number** | | **Node number** |
| --- | --- | --- | --- | --- | --- | --- | --- | --- | --- | --- | --- | --- | --- | --- | --- | --- | --- | --- | --- | --- | --- | --- | --- |
| **Early stolon number** | | | 0.646 | 0.111 | | 0.079 | | 0.058 | | 0.077 | | -0.120 | | 0.018 | | 0.016 | | -0.023 | | -0.048 | | -0.078 | |
| **Early stolon length** | | | 0.111 | 0.388 | | 0.044 | | 0.053 | | 0.033 | | -0.072 | | 0.009 | | 0.033 | | -0.022 | | -0.033 | | -0.044 | |
| **Corolla length** | | | 0.079 | 0.044 | | 0.661 | | 0.054 | | -0.139 | | -0.151 | | 0.066 | | 0.257 | | -0.099 | | -0.065 | | -0.158 | |
| **Corolla width** | | | 0.058 | 0.053 | | 0.054 | | 0.555 | | -0.085 | | 0.030 | | 0.059 | | 0.024 | | -0.026 | | -0.056 | | -0.030 | |
| **Days to flower** | | | 0.077 | 0.033 | | -0.139 | | -0.085 | | 0.757 | | 0.175 | | -0.094 | | -0.252 | | 0.061 | | 0.009 | | 0.129 | |
| **Late stolon number** | | | -0.120 | -0.072 | | -0.151 | | 0.030 | | 0.175 | | 1.131 | | 0.019 | | -0.452 | | 0.159 | | 0.090 | | 0.381 | |
| **Late stolon length** | | | 0.018 | 0.009 | | 0.066 | | 0.059 | | -0.094 | | 0.019 | | 0.400 | | 0.097 | | 0.023 | | 0.013 | | -0.022 | |
| **Stem thickness** | | | 0.016 | 0.033 | | 0.257 | | 0.024 | | -0.252 | | 0.452 | | 0.097 | | 1.096 | | -0.165 | | -0.076 | | -0.362 | |
| **Flower number** | | | -0.023 | -0.022 | | -0.099 | | -0.026 | | 0.061 | | 0.159 | | 0.023 | | -0.165 | | 0.940 | | 0.379 | | 0.160 | |
| **Branch number** | | | -0.048 | -0.033 | | -0.065 | | -0.056 | | 0.009 | | 0.090 | | 0.013 | | -0.076 | | 0.379 | | 0.704 | | 0.115 | |
| **Node number** | | | -0.078 | -0.044 | | -0.158 | | -0.030 | | 0.129 | | 0.381 | | -0.022 | | -0.362 | | 0.160 | | 0.115 | | 0.624 | |

|  |  | **Early stolon number** | | | **Early stolon length** | | **Corolla length** | | **Corolla width** | | **Days to flower** | | **Late stolon  number** | | **Late stolon  length** | | **Stem thickness** | | **Flower number** | | **Branch number** | | **Node number** |
| --- | --- | --- | --- | --- | --- | --- | --- | --- | --- | --- | --- | --- | --- | --- | --- | --- | --- | --- | --- | --- | --- | --- | --- |
| **Early stolon number** | | | 1 | 0.222 | | 0.123 | | 0.097 | | 0.121 | | -0.14 | | 0.036 | | 0.027 | | -0.03 | | -0.075 | | -0.124 | |
| **Early stolon length** | | | 0.222 | 1 | | 0.109 | | 0.144 | | 0.086 | | -0.062 | | 0.024 | | 0.086 | | -0.036 | | -0.053 | | -0.073 | |
| **Corolla length** | | | 0.123 | 0.109 | | 1 | | 0.09 | | -0.198 | | -0.242 | | 0.171 | | 0.318 | | -0.102 | | -0.095 | | -0.246 | |
| **Corolla width** | | | 0.097 | 0.144 | | 0.09 | | 1 | | -0.113 | | 0.027 | | 0.181 | | 0.031 | | -0.027 | | -0.063 | | -0.037 | |
| **Days to flower** | | | 0.121 | 0.086 | | -0.198 | | -0.113 | | 1 | | 0.154 | | -0.229 | | -0.234 | | 0.065 | | 0.012 | | 0.189 | |
| **Late stolon number** | | | -0.14 | -0.062 | | -0.242 | | 0.027 | | 0.154 | | 1 | | 0.03 | | -0.417 | | 0.173 | | 0.101 | | 0.406 | |
| **Late stolon length** | | | 0.036 | 0.024 | | 0.171 | | 0.181 | | -0.229 | | 0.03 | | 1 | | 0.146 | | 0.036 | | 0.021 | | -0.056 | |
| **Stem thickness** | | | 0.027 | 0.086 | | 0.318 | | 0.031 | | -0.234 | | -0.417 | | 0.146 | | 1 | | -0.175 | | -0.086 | | -0.436 | |
| **Flower number** | | | -0.03 | -0.036 | | -0.102 | | -0.027 | | 0.065 | | 0.173 | | 0.036 | | -0.175 | | 1 | | 0.452 | | 0.208 | |
| **Branch number** | | | -0.075 | -0.053 | | -0.095 | | -0.063 | | 0.012 | | 0.101 | | 0.021 | | -0.086 | | 0.452 | | 1 | | 0.155 | |
| **Node number** | | | -0.124 | -0.073 | | -0.246 | | -0.037 | | 0.189 | | 0.406 | | -0.056 | | -0.436 | | 0.208 | | 0.155 | | 1 | |

**Table S4.** Genetic correlation matrix for the 11 traits measured in the base generation (**G0**) on 828 *Mimulus guttatus* plants. Values represent genetic correlations (***rg***) estimated from the genetic variance-covariance (***G***) matrix obtained using Baysian mixed models. Positive values indicate traits that increase together, while negative values indicate opposing genetic associations.

**Table S5.** Summary of ANOVA testing for differences in early stolon number among control groups. Groups include the entire first generation, a resurrected ancestral population grown in generation 4, and the fourth-generation control line. Results show no significant differences in mean stolon number among groups (*P* = 0.107; see Figure 2C).

|  | df | Sum sq. | Mean sq. | *F* value | *P*-value |
| --- | --- | --- | --- | --- | --- |
| Control group | 2 | 7.2 | 3.622 | 2.238 | 0.1070 |
| Block | 1 | 4.9 | 4.867 | 3.008 | 0.0831 |
| Residuals | 1295 | 2095.2 | 1.618 |  |  |

**Table S6.** Pairwise comparisons for PC2 across groups in generation 4, corresponding to Figure 3D. A total of 10 pairwise comparisons were performed among generation 4 groups: the control (C), high (H1, H2), and low (L1, L2). *P*-values were adjusted for multiple comparisons using Tukey’s method. Bold values denote statistically significant differences (*P* < 0.05).

| Pairwise contrasts | Estimate | *P*-value |
| --- | --- | --- |
| C vs. H1 | -0.673 | **< 0.001** |
| C vs. H2 | -0.846 | **< 0.001** |
| C vs. L1 | 0.004 | 0.999 |
| C vs. L2 | 0.162 | 0.999 |
| H1 vs. H2 | -0.173 | 0.805 |
| H1 vs. L1 | -0.676 | **< 0.001** |
| H1 vs. L2 | -0.689 | **< 0.001** |
| H2 vs. L1 | -0.850 | **< 0.001** |
| H2 vs. L2 | -0.862 | **< 0.001** |
| L1 vs. L2 | 0.013 | 0.999 |

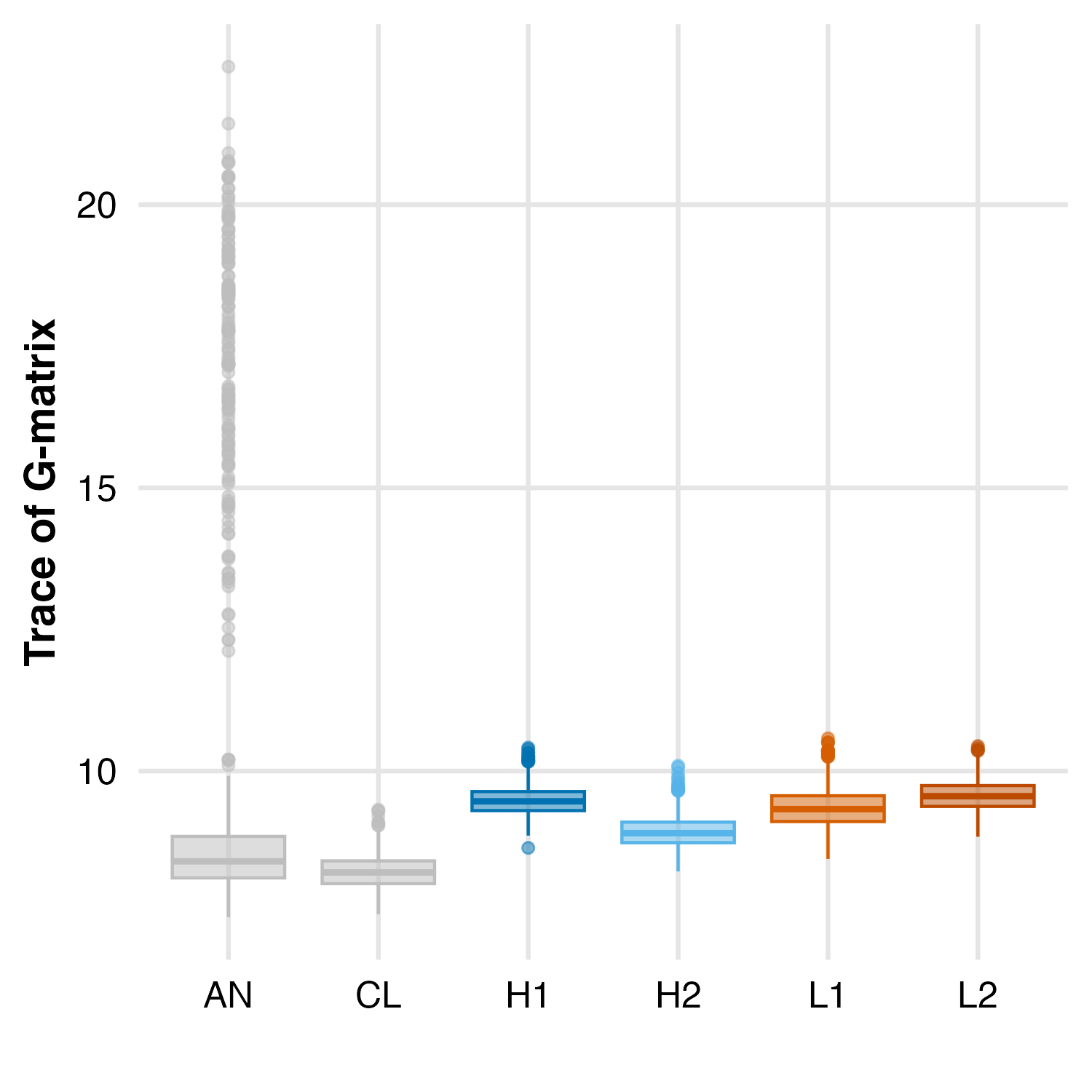

**Figure S4.** Boxplots showing the distributions of bootstrapped **G** matrix sizes (trace of the **G** matrix; sum of eigenvalues) across the ancestral population, control, and selection lines in generation 4. Statistical comparisons using Analysis of Variance (ANOVA) are shown in Table S6.

**Table S7.** Pairwise comparisons of total genetic variance (trace) across groups in generation 4. A total of 15 pairwise comparisons were performed among generation 4 groups: the ancestral population (AN), control (C), high (H1, H2), and low (L1, L2). *P*-values were adjusted for multiple comparisons using Tukey’s method. Bold values denote statistically significant differences (*P* < 0.05).

| Pairwise contrasts | Estimate | *P*-value |
| --- | --- | --- |
| AN vs. C | 1.45 | **< 0.001** |
| AN vs. H1 | 0.20 | **0.011** |
| AN vs. H2 | 0.74 | **< 0.001** |
| AN vs. L1 | 0.33 | **< 0.001** |
| AN vs. L2 | 0.12 | 0.373 |
| C vs. H1 | -1.25 | **< 0.001** |
| C vs. H2 | -0.71 | **< 0.001** |
| C vs. L1 | -1.12 | **< 0.001** |
| C vs. L2 | -1.33 | **< 0.001** |
| H1 vs. H2 | 0.54 | **< 0.001** |
| H1 vs. L1 | 0.13 | 0.274 |
| H1 vs. L2 | -0.08 | 0.736 |
| H2 vs. L1 | -0.41 | **< 0.001** |
| H2 vs. L2 | -0.62 | **< 0.001** |
| L1 vs. L2 | -0.21 | **0.005** |

**Table S8.** Pairwise comparisons of Krzanowski’s common subspaces across groups in generation 4. A total of 15 pairwise comparisons were performed among generation 4 groups: the ancestral population (AN), control (C), high (H1, H2), and low (L1, L2). *P*-values were adjusted for multiple comparisons using Tukey’s method. Bold values denote statistically significant differences (*P* < 0.05)

| Pairwise contrasts | Estimate | *P*-value |
| --- | --- | --- |
| AN vs. C | 0.13 | **< 0.001** |
| AN vs. H1 | 0.22 | **< 0.001** |
| AN vs. H2 | 0.24 | **< 0.001** |
| AN vs. L1 | 0.17 | **< 0.001** |
| AN vs. L2 | 0.15 | **< 0.001** |
| C vs. H1 | 0.10 | **< 0.001** |
| C vs. H2 | 0.11 | **< 0.001** |
| C vs. L1 | 0.04 | 0.366 |
| C vs. L2 | 0.02 | 0.924 |
| H1 vs. H2 | 0.02 | 0.961 |
| H1 vs. L1 | -0.06 | 0.039 |
| H1 vs. L2 | -0.08 | 0.001 |
| H2 vs. L1 | -0.07 | **0.002** |
| H2 vs. L2 | -0.09 | **< 0.001** |
| L1 vs. L2 | -0.02 | 0.924 |
